# Monocyte Adhesion and Transmigration Through Endothelium Following Cardiopulmonary Bypass Shearing is Mediated by IL-8 Signaling

**DOI:** 10.1101/2023.06.05.543811

**Authors:** Hao Zhou, Lan N Tu, Cecilia Giachelli, Vishal Nigam, Marta Scatena

**Affiliations:** University of Washington, Seattle, WA; Seattle Children’s Hospital, Seattle, WA

**Author notes:** Corresponding author: Marta Scatena, PhD, Department of Bioengineering University of Washington, Box 358056, Seattle, WA 98195.

## Abstract

**Background:** The use of cardiopulmonary bypass (CPB) can induce sterile systemic inflammation that contributes to morbidity and mortality, especially in children. Patients have been found to have increased expression of cytokines and transmigration of leukocytes during and after CPB. Previous work has demonstrated that the supraphysiologic shear stresses present during CPB are sufficient to induce proinflammatory behavior in non-adherent monocytes. The interactions between shear stimulated monocytes and vascular endothelial cells have not been well studied and have important translational implications.

**Methods:** To test the hypothesis that non-physiological shear stress experienced by monocytes during CPB affects the integrity and function of the endothelial monolayer via IL-8 signaling pathway, we have used an in vitro CPB model to study the interaction between THP-1 monocyte-like cells and human neonatal dermal microvascular endothelial cells (HNDMVECs). THP-1 cells were sheared in polyvinyl chloride (PVC) tubing at 2.1 Pa, twice of physiological shear stress, for 2 hours. Interactions between THP-1 cells and HNDMVECs were characterized after coculture.

**Results:** We found that sheared THP-1 cells adhered to and transmigrated through the HNDMVEC monolayer more readily than static controls. When co-culturing, sheared THP-1 cells also disrupted in the VE-cadherin and led to reorganization of cytoskeletal F-actin of HNDMVECs. Treating HNDMVECs with IL-8 resulted in upregulation of vascular cell adhesion molecule 1 (VCAM-1) and intercellular adhesion molecule 1 (ICAM-1) while also increasing the adherence of non-sheared THP-1 cells. Preincubating HNDMVECs with Reparixin, an inhibitor of CXCR2/IL-8 receptor inhibited sheared THP-1 cell adhesion to the HNDMVECs.

**Conclusions:** These results suggested that IL-8 not only increases the endothelium permeability during monocyte migration, but also affects the initial adhesion of monocytes in a CPB setup. This study revealed a novel mechanism of post-CPB inflammation and will contribute to the development of targeted therapeutics to prevent and repair the damage to neonatal patients.

**Highlights:**

- Shear stress in a CPB-like environment promoted the adhesion and transmigration of monocytes to and through endothelial monolayer.
- Treating endothelial monolayer with sheared monocytes led to disruption of VE-cadherin and reorganization of F-actin.
- Interaction between sheared monocytes resulted in a significant increase of IL-8 release.
- Inhibiting IL-8 receptor prevented sheared monocyte adhesion, while IL-8 promoted naive monocyte adhesion.

## Introduction

Cardiopulmonary bypass (CPB) is a technique that maintains the blood circulation in an extracorporeal circuit, in which blood is propelled by mechanical pumps and flows in plastic tubing. This technique provides the surgeon with a bloodless field to conduct open-heart surgery, while minimizing the ischemic damage to the organs. However, since the blood components are exposed to an environment that is very different from the native blood vessel during CPB, the incidence of complication occurs at a rate of 29.2% [1]. These complications can lead to systemic inflammation and multiorgan dysfunction, which translates to a mortality rate of 10.7% in the pediatric patients. Although modifications have been made to suppress the inflammatory response through corticosteroid administration, coating the CPB tubing, and filtrating the blood postoperatively, no significant improvement has been reported [2–6]. At present, finding a CPB-specific pathway that leads to the inflammatory response in pediatric patients is the key to reducing the complication and motility rate.

Post-CPB complications are largely associated with the secretion of inflammatory cytokines, such as Interleukin (IL)-1β, IL-6, IL-8 and Tumor necrosis factor alpha (TNF-α) during and after bypass [7–16] and disruption of the endothelial barrier function [17,18]. However, most current studies were based on clinical samples, which are unlikely to decouple the effects of the CPB and of the surgical intervention. Therefore, it remains challenging to find targeted pathways that prevent the inflammatory response specific to CPB. Given that blood cells can experience three times more shear stress in a CPB circuit compared to that in the native blood vessel [19], studying the correlation between CPB shear and cytokine released by cells can help understand CPB-specific pathways. Recently, we developed an *in vitro* model that was designed to distinguish the non-CPB factors such as surgical trauma, blood transfusion, drug administration, and the specific condition of the patient from CPB-specific inflammatory changes [19]. Using this model, we demonstrated that CPB shear stress specifically upregulated the expression of IL-8 and TNF-α in monocytes, as well as giving rise to a subpopulation of monocytes undergoing TNF-α-mediated necroptosis. While TNF-α-mediated inflammatory response of cells in the CPB setting has been well-studied [19–21], the role of IL-8 in regulating the cellular response after CPB is still unclear.

IL-8 is a chemoattractant that regulates the recruitment of neutrophils, basophils, and T-cell during inflammation. Cells that can secrete IL-8 include monocytes, macrophages, neutrophils, and endothelial cells. The receptors for IL-8, CXC chemokine receptor (CXCR)1 and CXCR2, are also reported to be present on a wide range of cell types, including endothelial cells, monocytes, neutrophils, fibroblasts, with the former having more specific affinity to the IL-8 than the latter [22]. Evidence has shown that in CPB patients, perioperative plasma IL-8 level is positively related to the length of inotropic support, length of mechanical ventilation, and incidence of acute kidney injuries [23–27]. Thus, IL-8-mediated signaling pathways may play an important role in the onset of CPB-induced inflammatory response at the cellular level. In this study we used a novel in vitro CPB model to characterize the molecular mechanism of how leukocytes affected by CPB shear can induce inflammatory response in endothelial cells. We identified IL-8/CXCR2 signaling pathway as a regulator of leukocyte adhesion and transmigration through the endothelial cell monolayer.

## Methods

The data that support the findings of this study are available from the corresponding author upon reasonable request.

### Cell line and culture methods

Primary human neonatal dermal microvascular cells (HNDMVECs, CC-2516) were purchased from Lonza (Walkersville, MD) and cultured in endothelial cell growth medium MV2 (ECG MV2, C-22121) (PromoCell, Heidelberg, Germany) and 100U/mL penicillin-streptomycin (Gibco, Waltham, MA). A HEK293T cell line was purchased from Invitrogen (Waltham, MA) and cultured in Dulbecco’s Modified Eagle Medium (DMEM) (Gibco, Waltham, MA) with 10% fetal bovine serum (FBS) (Atlanta Biologicals, Flowery Branch, GA) and 100U/mL penicillin-streptomycin. The medium was changed every 2 days and the cells were harvested by trypsinization. The human acute leukemia monocytic cell line THP-1 (TIB-202) was purchased from American Type Culture Collection (ATCC, Manassas, VA) and cultured in Roswell Park Memorial Institute (RPMI) 1640 medium (ATCC Modification) (Gibco, Waltham, MA) with 10% FBS. The medium was changed every 2 days. Cells were rinsed with fresh medium twice after pretreatment.

### Lentiviral production and transduction

A THP-1 cell line that expresses GFP (G-THP-1) was generated though lentiviral transduction. Vectors that contain plasmids of pSL3 (vesicular stomatitis virus G envelope), pSL4 (HIV-1 gag/pol packing genes), pSL5 (rev gene required for HIV-1 envelope protein expression) were a gift from Dr. Murry (University of Washington, Seattle, WA). The plasmid vector pCDH-EF1-MCS-T2A-copGFP was purchased from System Biosciences. The lentiviral vector was packaged in HEK293T cells as previously described. Briefly, 3x106 of HEK293T cells were seeded in 10-cm dishes to reach 60% confluence in DMEM with 10% FBS and 100U/mL penicillin-streptomycin. The culture media was changed prior to transfection. A total of 20ug plasmid DNA (7.5 ug pCDH-EF1-MCS-T2A-copGFP, 2.5 ug pSL3, 6.7 ug pSL4, 3.3 ug pSL5) and 24 uL of lipofectamine 2000 dissolved in 1600 uL Opti-MEM were used for the transfection of one dish. The media was replaced after incubating for 6 hours at 37°C with 5% CO2. The virus supernatant was collected at 48 hours from the media and filtered through a 0.45-µm filter. The media containing the virus was applied to the THP-1 cells at a multiplicity of infection of 6 while spinning at 800xg for 30 minutes. The transduced THP-1 cells (G-THP-1) were allowed to expand and sorted for GFP expression using a to obtain over 92% transduction efficiency.

### THP-1 Adhesion Assay

HNDMVECs were seeded on 24-well plates at 5000 cells/cm^2^ and cultured 7 days to form a confluent monolayer. Monocytes were sheared as previously described. Briefly, G-THP-1 cells at a density of 2 million cells/mL were sheared in a 10 foot-long Masterflex Tygon E-3603 L/S13 pump tubing (Cole-Parmer), using a Masterflex miniflex pump model 115/230 VAC 07525-20 (Cole-Parmer) at 10 mL/min for 2 hours. The tubing was mostly submerged in a water bath with temperature control at 37C. At the end of shear, the G-THP-1 cells were collected, spun down and resuspended in fresh media. As a control, G-THP-1 cells were incubated in a polyvinylchloride (PVC) flask at a density of 2 million cells/mL for 2 hours. For the pretreatment groups, HNDMVECs were either incubated with 5 nM of Reparixin (Sigma-Aldrich) for 30 minutes or 200 ng/mL of IL-8 for 1 hour. The sheared and statically incubated G-THP-1 cells were then added to a monolayer of HNDMVECs (1x10^6^ cells per well), followed by incubation for 1 hour. Nonadherent THP-1 cells were gently washed twice with sterile phosphate-buffered saline (PBS). The adhered monocytes were visualized using a Nikon Eclipse TE200 microscope (Tokyo, Japan) and a Photometric CoolSNAP MYO camera (Tucson, AZ). The experiments were run in triplicates. The number of adherent G-THP-1 cells were counted in five randomly selected visible fields and quantified using ImageJ [28].

### THP-1 Transmigration Assay

HNDMVECs were seeded in the upper chamber of a transwell tissue culture insert (6.5 mm diameter, 8 μm pore size polycarbonate membrane; Corning, NY) at 5000 cells/cm^2^ and cultured 7 days to form a confluent monolayer. Sheared and statically incubated G-THP-1 cells were then added to the upper chamber (2x10^5^ cells/well) with the lower chamber filled with RPMI 1640 media. After 24 hours of incubation, the upper chamber was removed and the THP-1 cells in the lower chamber were visualized using a Nikon Eclipse TE200 microscope and a Photometric CoolSNAP MYO camera and quantified with ImageJ. The ability of transendothelial migration was determined by counting the migrated G-THP-1 cells in the lower chamber [29]. The experiment was run in triplicates. The number of adherent G-THP-1 cells were counted in five randomly selected visible fields and quantified using ImageJ [28].

### Immunofluorescent Staining

HNDMVECs were seeded on 8-chamber slides (354118, Corning, NY) at 5000 cells/cm^2^ and cultured 7 days to form a confluent monolayer. The sheared or statically incubated G-THP-1 cells were then added to a monolayer of HNDMVECs (1x10^6^ cells per well), followed by incubation for 6 hours. The cell cocultures were washed twice with sterile PBS, fixed with formalin. For staining, the fixed cells were permeabilized with 0.1% Triton-X-100 for 10 minutes, blocked with 1% BSA and human BD Fc block (BD Bioscience, Franklin Lakes, NJ) for 1 hour, followed by incubation with 1:200 vascular endothelial cadherin (VE-cadherin) monoclonal antibody (16B1), Biotin (Thermofisher) for 1 hour, 1:500 streptavidin, Alexa Fluor 594 Conjugate (Thermofisher) for 45 minutes. Nuclei were stained with DAPI. The slides were then washed, mounted, and imaged using a Leica DMI6000 microscope with Leica SP8X confocal. VE-cadherin staining was quantified using ImageJ [28].

### Rhodamine-Phalloidin Labeling

HNDMVECs were seeded on 8-chamber slides at 5000 cells/cm^2^ and cultured 7 days to form a confluent monolayer. The sheared or statically incubated G-THP-1 cells were then added to a monolayer of HNDMVECs (1x10^6^ cells per well), respectively, followed by incubation for 6 hours. The cell cocultures were washed twice with sterile PBS, fixed with formalin, permeabilized with 0.1% Triton-X-100, and blocked with 1% BSA. The cells were then incubated with Rhodamine Phalloidin (ThermoFisher, R415) for 20 minutes at room temperature, stained with DAPI, washed, mounted, and imaged using a Leica DMI6000 microscope with Leica SP8X confocal.

### Enzyme-linked Immunosorbent Assays (ELISA)

The media supernatants of the HNDMVECs and sheared or statically incubated THP-1 cells were collected at 0.5, 1, 3, and 6 hours into the coculture. The levels of IL-1β, IL-6, IL-8 and TNF-α were measured using ELISA kits from Invitrogen (Catalog# BMS224-2, 88-7066-88, 88-8086-88, BMS223-4). Experiments were run in both biological and technical triplicates.

### Quantitative Real-time PCR

HNDMVECs were treated with human recombinant IL-8 for 1 hour, and total RNA was isolated using the RNeasy Mini Kit (Qiagen) and RNase-Free DNAse Set (Qiagen). Total RNA (150 ng) was used for cDNA synthesis using the Omniscript RT Kit (Qiagen). Quantitative real-time PCR was performed on a QuantStudio 6 Pro real-time PCR system and Taqman Universal PCR Master Mix according to the manufacturer instructions. Both biological and technical triplicates were run. Taqman primers of selectin E (Hs00174057_m1), intercellular adhesion molecule 1 (ICAM1) (Hs00164932_m1) and vascular cell adhesion molecule 1 (VCAM1) (Hs01003372_m1) were purchased from Thermofisher.

### Statistical Analysis

All experiments were run in triplicates. All quantitative data were expressed as mean ± standard deviation within groups. Pairwise comparisons between groups were conducted using ANOVA test and Tukey’s post-hoc test. Statistical significance is denoted by ‘*’. P values less than 0.05 are indicated by single symbol and P values less than 0.01 are indicated by double symbols.

## Results

### 1. CPB shear promotes THP-1 adhesion to and transmigration through endothelial monolayer

Monocytes infiltration is commonly observed in the inflammatory response in CPB [30–32]. During this process, activated monocytes adhere to the vascular wall, transmigrate through the intercellular tight junctions, and reach different organs [33–36]. To characterize the response of monocytes to specifically to CPB shear, we sheared the THP-1 cells in the CPB circuit, cocultured them with confluent HNDMVECs for 1 hour and quantified the number of attached THP-1 cells. As shown in Figure 1A, exposing THP-1 to shear stress resulted in a two-fold increase in THP-1 cells adhering to the HNDMVEC monolayer. Furthermore, significantly more sheared THP-1 cells were observed in the bottom chamber of the transwells that had been seeded with HNDMVECs compared with static THP-1 cells (Figure 1D). These results suggest that exposure to CPB shear induced monocytic cells to be more adherent to and likely to transmigrate through a layer of endothelial cells.

**Figure 1.**
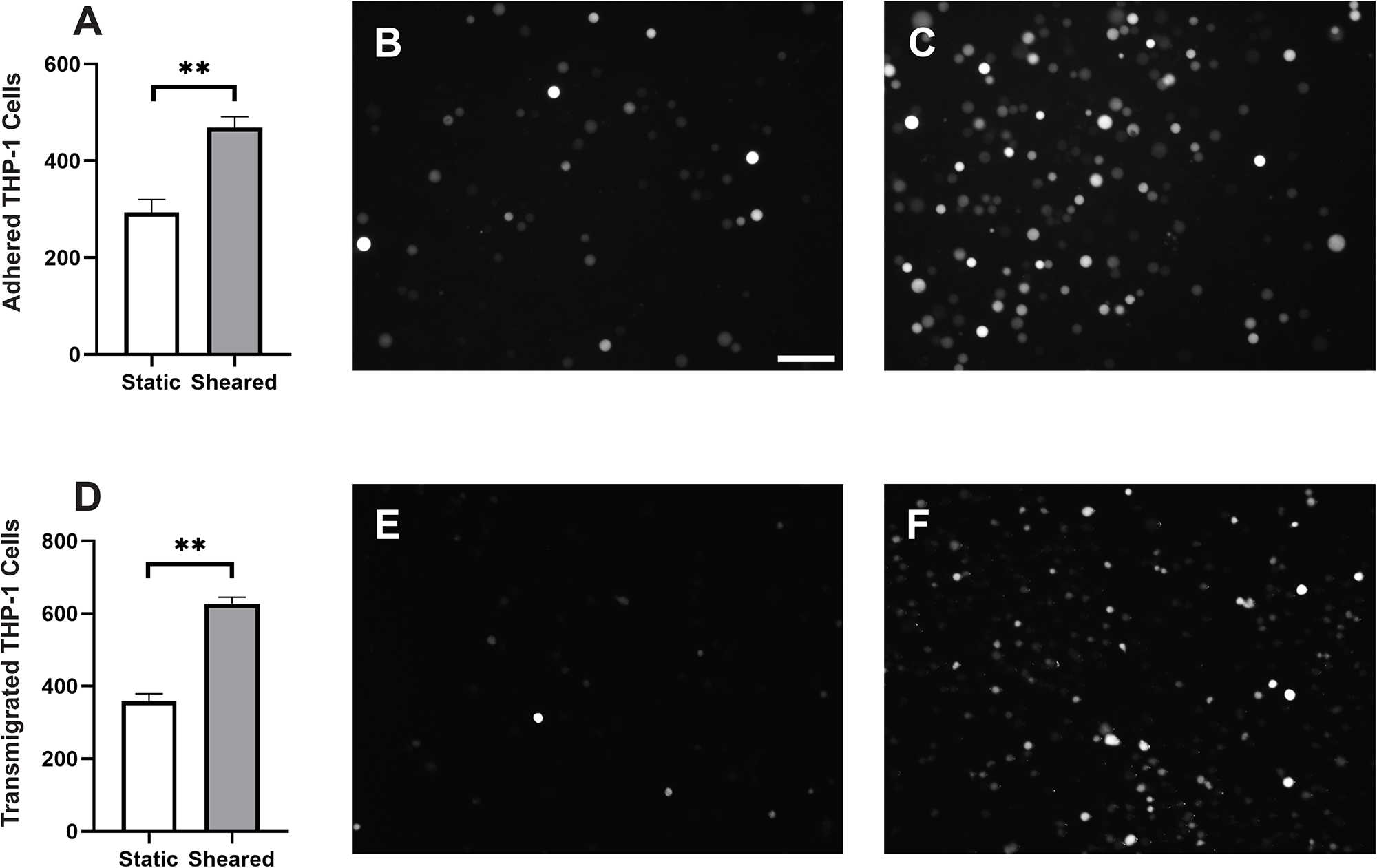
A) Quantitative analysis of adhered THP-1 cells on the endothelial cell monolayer. B) Adhesion of THP-1 cells incubated statically in a PVC flask. C) Adhesion of THP-1 cells sheared in a CPB circuit. D) Quantitative analysis of transmigrated THP-1 cells through endothelial cell monolayer. E) Transmigrated THP-1 cells incubated statically in a PVC flask. F) Transmigration of THP-1 cells sheared in a CPB circuit. Scale bar = 100 μm. Additional images see Figure S1. Statistical significance is denoted by ‘*’. P values less than 0.05 are indicated by single symbol and P values less than 0.01 are indicated by double symbols.

### 2. Treating confluent HNDMVECs with sheared THP-1 cells results in disrupted intercellular junction

The increasing transmigration behavior of sheared monocytes prompted us to examine if the sheared cells were disrupting the barrier function in the endothelial monolayer. To examine the intercellular adherens junction and reorganization of cytoskeleton, HNDMVECs were stained for VE-cadherin and F-actin after co-culture with sheared or static THP-1 cells. Co-culturing with sheared THP-1 cells resulted in the loss of adherens junction while the tight junction stayed intact in endothelial cells cultured with static THP-1s (Figure 2A, C). Similarly, the static group showed robust actin filament lining intracellularly (Figure 2B, D). In the sheared group, we observed THP-1 cells clustered at the intercellular area and the F-actin filament was reorganized to compromise the space (Figure 3A, C), which implies increasing motility of the endothelial cells.

**Figure 2.**
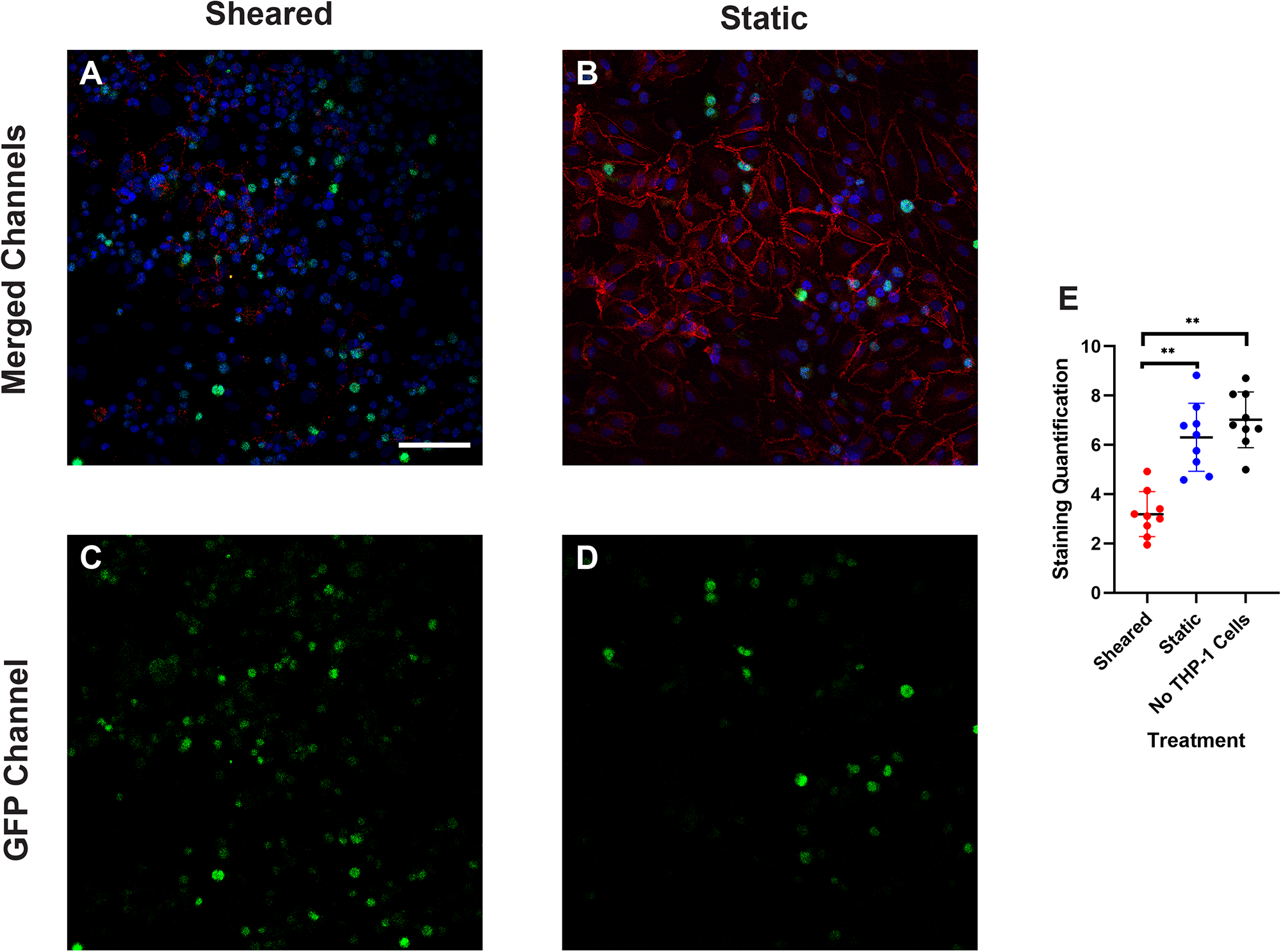
VE-cadherin of HNDMVECs cocultured with A) sheared THP-1 cells and B) THP-1 cells statically incubated in PVC flask, visualized by immunofluorescent staining. C) and D) show the locations of GFP positive THP-1 cells in A) and B) respectively. Additional images see Figure S2. Scale bar = 100 μm.

**Figure 3.**
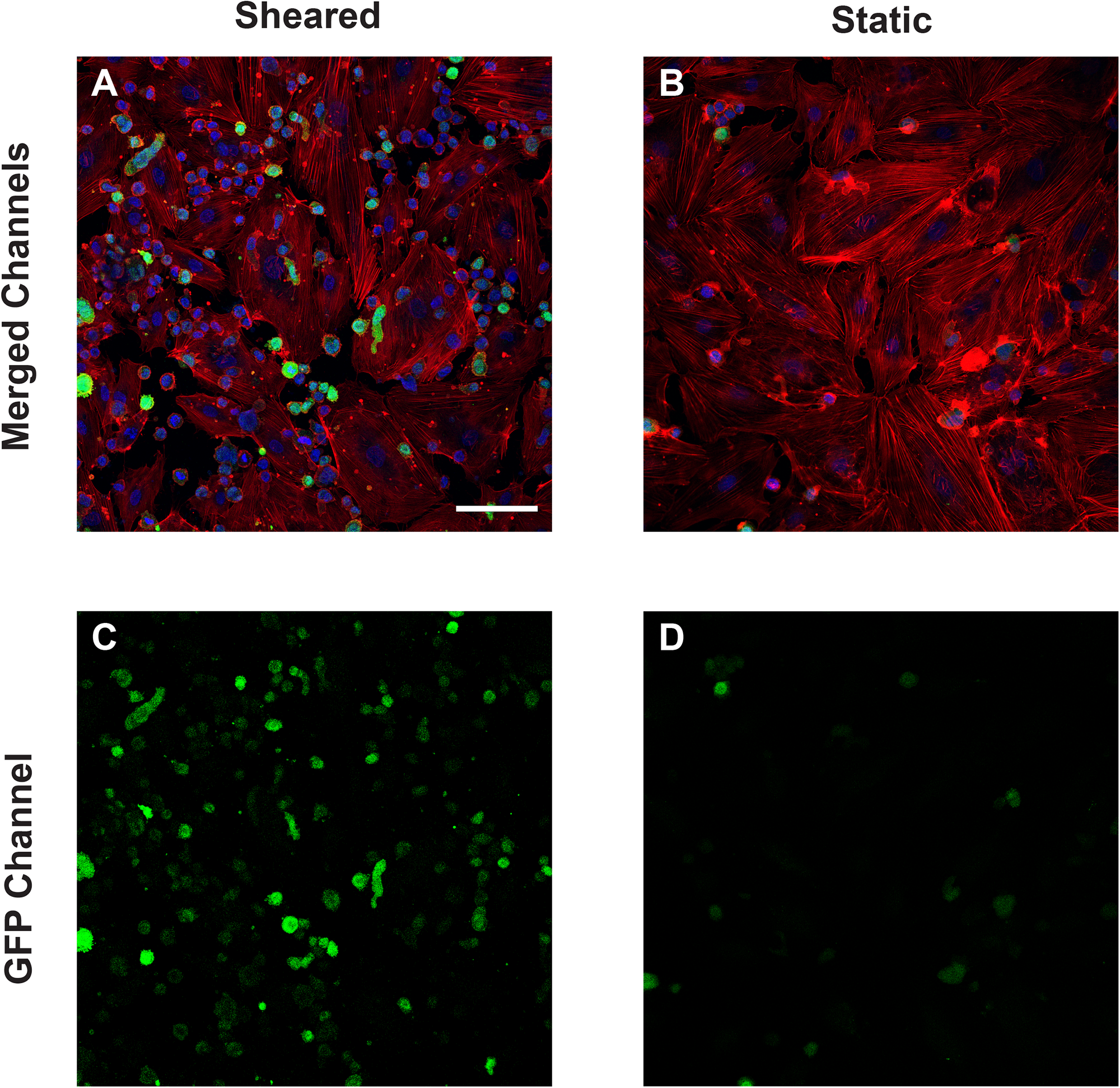
F-actin of HNDMVECs cocultured with A) sheared THP-1 cells and B) THP-1 cells statically incubated in PVC flask, stained by rhodamine phalloidin. C) and D) show the locations of GFP positive THP-1 cells in A) and B) respectively. Additional images see Figure S3. Scale bar = 100 μm.

### 3. Interaction between sheared THP-1 and HNDMVECs results in increased IL-8 release level

To analyze the inflammatory cytokines secreted by sheared or static THP-1 cells and HNDMVECs, we collected coculture media at 0.5, 1, 3, and 6 hours and measured the levels of IL-1β, IL-6, IL-8 and TNF-α using ELISA. Sheared group released significantly more IL-8 than the static group at every time point (Figure 4A). The level of IL-8 increased over the period of 6 hours in the shear group while it leveled off at 3 hours in the static group (Figure 4A). IL-6 level in the sheared group significantly increased at 3 and 6 hours but the concentration was about 20-fold lower than IL-8 (Figure 4B). No significant release of IL-1β and TNF-α were detected throughout the coculture (Figure 4C, TNF-α data not shown). These findings indicate that treating HNDMVECs with sheared THP-1 specifically induced increasing IL-8 release.

**Figure 4.**
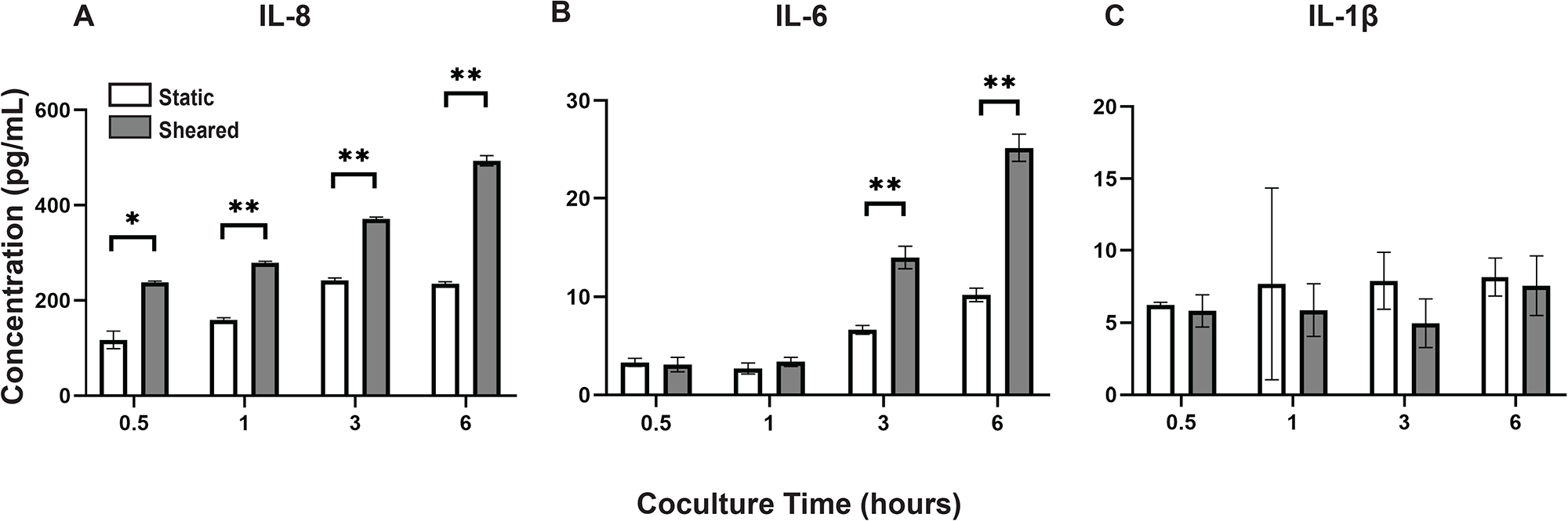
Cytokine levels of A) IL-8, B) IL-6, and C) IL-1β in the coculture media of sheared or static THP-1 cells and HNDMVECs measured at 0.5, 1, 3, and 6 hours. Statistical significance is denoted by ‘*’. P values less than 0.05 are indicated by single symbol and P values less than 0.01 are indicated by double symbols.

### 4. Reparixin inhibits the adhesion of sheared THP-1 cells while IL-8 promotes the adhesion of healthy THP-1 cells

Our results suggested that the adhesion of THP-1 cells on the endothelial cell is positively correlated to the IL-8 release during coculture. To further investigate the role of IL-8 in mediating the interaction between THP-1 cells and the endothelial cells, we treated the endothelial cells with Reparixin, an inhibitor of the IL-8 receptors, CXCR2, and then co-cultured with sheared THP-1 cells (CXCR2 staining of HNDMVECs see Figure S6). The Reparixin treatment group reduced THP-1 adhesion like the static group (Figure 5C). When we pre-treated HNDMVECs with 200 ng/ml IL-8, the adhesion of static THP-1 cells was promoted (Figure 6C, D). These results suggested that blocking IL-8 signaling through the CXCRs prevented the adhesion of shear-activated THP-1 cells.

**Figure 5.**
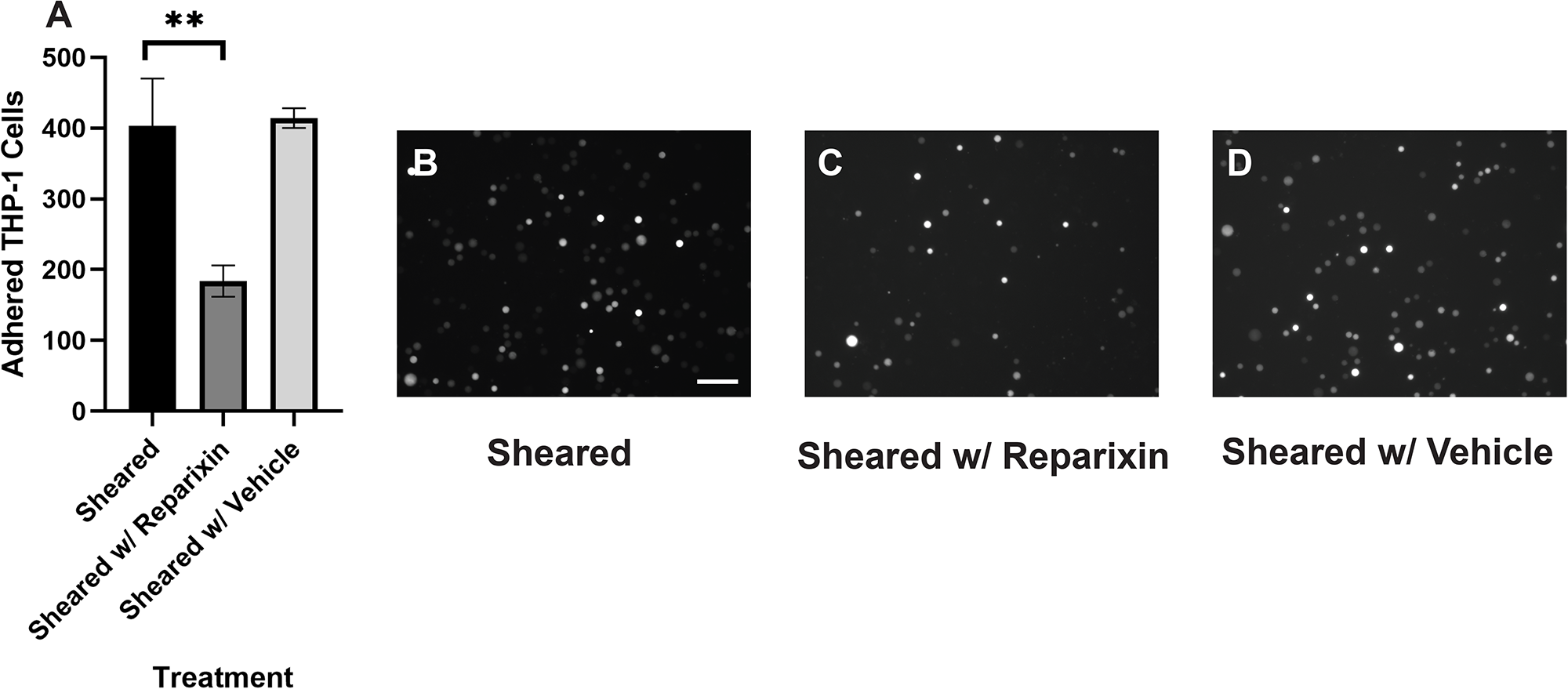
A) Quantitative analysis of sheared THP-1 cells adhering to B) untreated HNDMVECs, C) HNDMVECs pretreated with 5nM reparixin for 30 minutes, and D) HNDMVECs pretreated with PBS. Scale bar = 100 μm. Additional images see Figure S4. Statistical significance is denoted by ‘*’. P values less than 0.05 are indicated by single symbol and P values less than 0.01 are indicated by double symbols.

**Figure 6.**
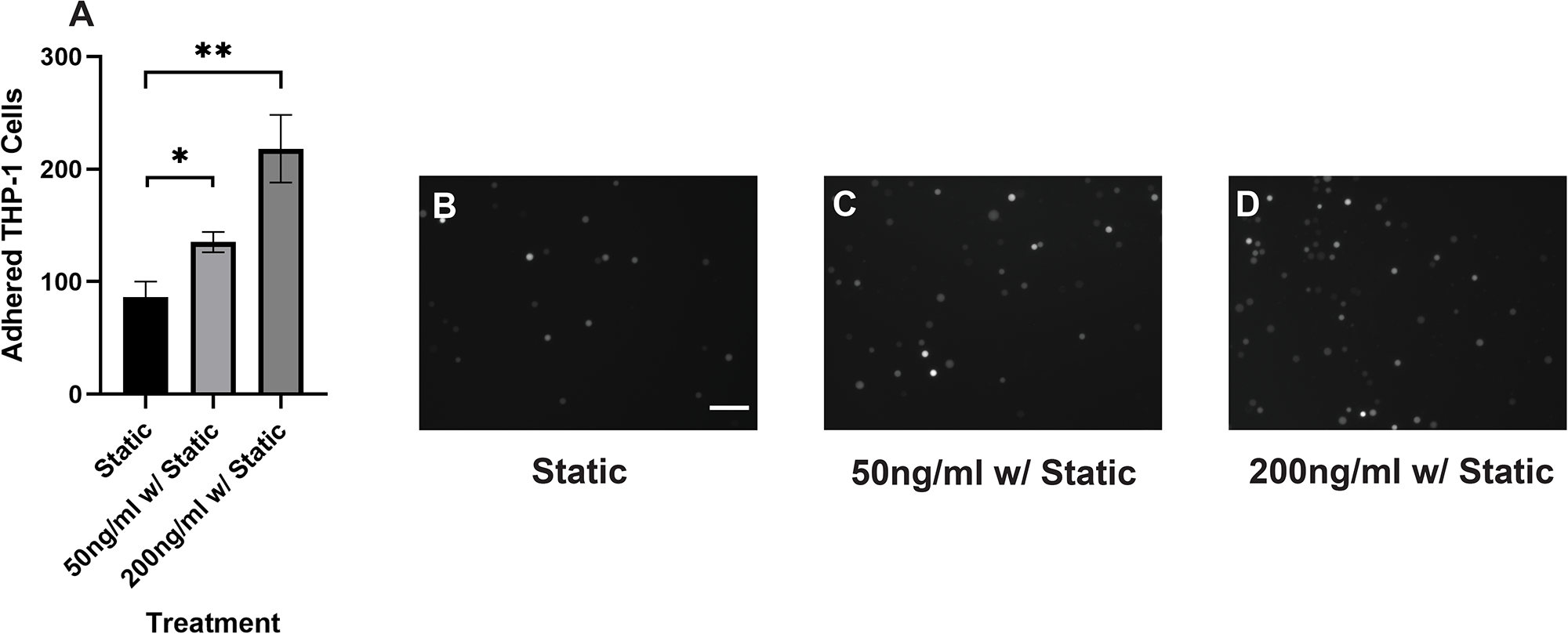
A) Quantitative analysis of statically incubated THP-1 cells adhering to B) untreated HNDMVECs, C) HNDMVECs pretreated with 50 ng/mL IL-8 for 1 hour, and D) HNDMVECs pretreated with 200 ng/mL IL-8 for 1 hour. Additional images see Figure S5. Scale bar = 100 μm. Statistical significance is denoted by ‘*’. P values less than 0.05 are indicated by single symbol and P values less than 0.01 are indicated by double symbols.

### 5. IL-8 upregulated adhesion molecules on the endothelial cells

In order to gain mechanistic insight into how IL-8 promotes myeloid cell adherence to endothelial cells, we examined the expression of adherence molecules in the endothelial cells. qPCR was performed to examine the expression of adhesion molecules, including e-selectin, VCAM1, and ICAM1. HNDMVECs were treated with IL-8 with TNF-α treatment being a positive control to stimulate the endothelial cells. All three adhesion molecules were upregulated in the positive control group (Figure 7). In the IL-8 group, there was a 5-fold increase in both the expressions of ICAM1 and VCAM1, while the expression of e-selectin remained unchanged (Figure 7). These results indicate that the increasing adhesion of sheared THP-1 cells was likely to be mediated by IL-8 signaling pathway through upregulated adhesion molecule expression on the endothelial cells.

**Figure 7.**
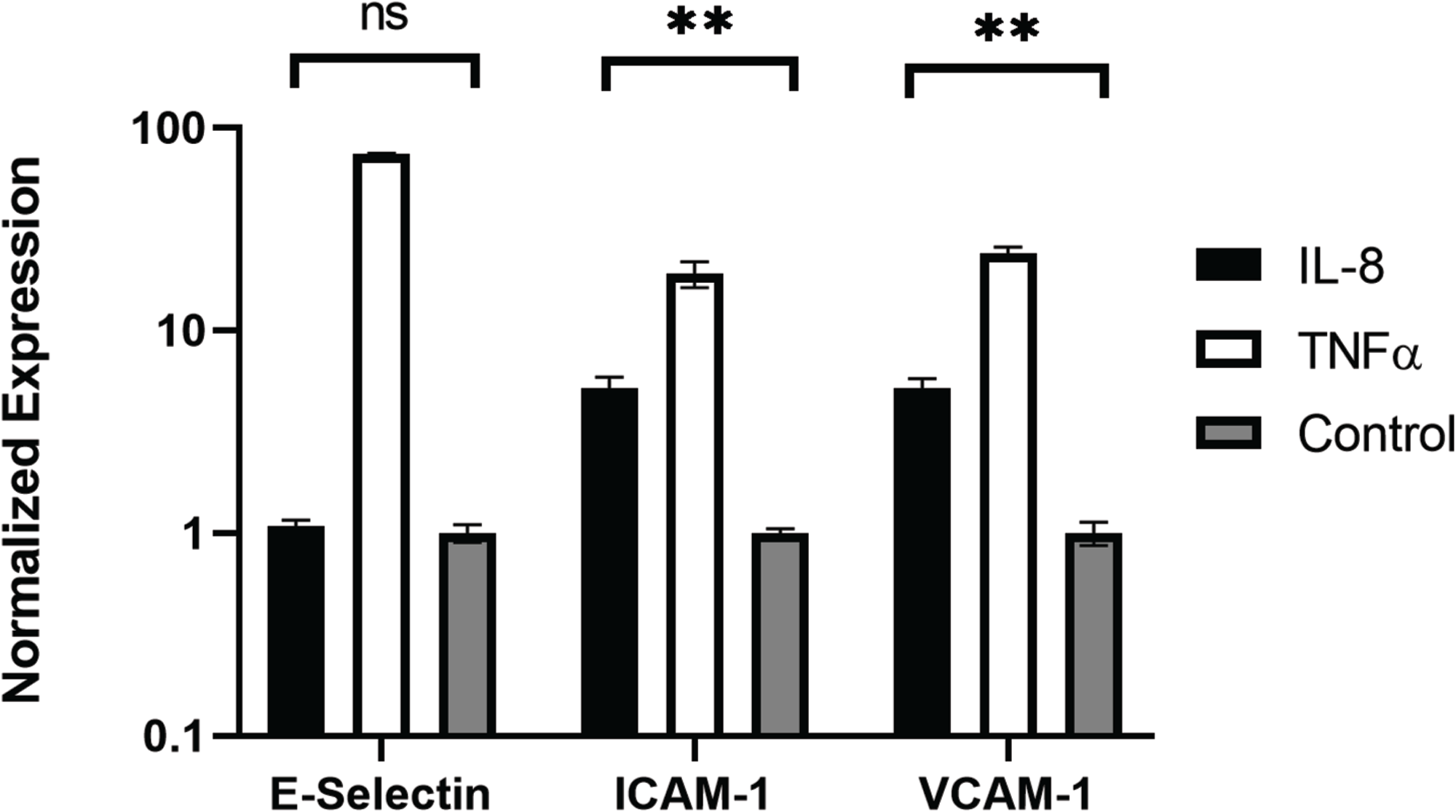
Expression of e-selectin, ICAM1, and VCAM1 in HNDMVECs treated with IL-8, TNF-α, and control measured by RT-qPCR. Statistical significance is denoted by ‘*’. P values less than 0.05 are indicated by single symbol and P values less than 0.01 are indicated by double symbols.

## Discussion

With this study to our knowledge, we are the first to show that CPB shear stress activated monocytes adhere to and migrate through an intact and unstimulated endothelial monolayer. We used an *in vitro* CPB model to probe the interaction between shear stress activated THP-1 monocytes and quiescent confluent HNDMECs. Our data demonstrated that an average CPB shear stress of 2.1 Pa or 21 dyne/cm^2^ promoted the adhesion and transmigration of monocytes to and through the endothelial monolayer. This was accompanied by the disruption of *adherens junction* and reorganization of cytoskeleton in the endothelial cells. IL-8, which was significantly upregulated in the coculture media, played a pivotal role in mediating the crosstalk between monocytes and the endothelial cells. Inhibiting the binding of IL-8 to the CXCRs on endothelial cells prevented sheared monocytes from adhering. Preincubating endothelial cells with IL-8 resulted in increasing adhesion of untreated monocytes, suggesting endothelial cells activation. Indeed, we found that treating with IL-8 induced ICAM1 and VCAM1 expression in HNDMVECs. This suggests that IL-8 may promote the adhesion of the CPB shear stress treated monocytes by activating a subset of adhesion molecules on the microvascular endothelial cells.

The sequela of leukocyte rolling and binding to activated endothelial cells and transmigration through the endothelial monolayer is a fundamental process of inflammation responding to mediators produced by a traumatic injury. We have recently shown that using *in vitro* CPB set up recapitulating a prolonged CPB surgery, monocytes circulating for 2 hours at 21 dyne/cm^2^ showed increased expression of IL-8 and TNF-α [19]. Now we also find that these cells can adhere to and transmigrate through an endothelial monolayer without additional stimulation. These results suggest that CPB shear stress is sufficient to promote monocyte adhesion and transmigration. Additionally, loss of endothelial membrane VE-cadherin staining correlated with formation of gaps in the monolayer. Importantly, these effects seem to be independent of the THP-1 interaction with the CPB circuit material, PVC, since control cells incubated statically in PVC containers showed much reduced adhesion and were comparable to cell statically cultured on tissue culture plastic. Interestingly, Baratchi et al., using a microfluidic system modeling the change in shear stress observed in aortic stenosis, recently described calcium flux dependent activation of monocytes in response to elevated shear stress and their ability to adhere to TNF-α stimulated HUVECs [37]. Our data strongly suggest that TNF-α was not playing a role in our CPB shear stress induced THP-1/endothelial cell interactions, as we did not find changes of TNF-α levels in the coculture medium, even if TNF-α was induced by CPB shear stress in monocytes as shown previously [19]. A potential explanation for the absence of TNF-α in the coculture is that CPB shear stress activated THP-1 media was refreshed before coculture with the endothelial monolayer, suggesting that when the shear stress stimulus is removed, the monocytes stop producing TNF-α. These results point to different factors mediating the monocyte-endothelial cell interaction in our system.

We show that the monocyte-endothelial cell interaction was promoted by addition of IL-8 and inhibited by the CXCR antagonist Reparixin. These results suggest that IL-8 mediates the interaction between monocytes and endothelial cells in CPB shear stress conditions. Accordingly, we have previously shown that IL-8 in the monocytes is directly upregulated by CPB shear stress [19]. Here we find that IL-8 was also elevated in the media of CPB shear stress stimulated THP-1 and endothelial cell cocultures. The results of the current study revealed that IL-8 is a main cytokine mediating the interaction between monocytes and endothelial cells in a CPB setting. Current study design limitation did not permit us to identify the cell type contributing to IL-8 release, however, both monocytes and endothelial cells have been shown to secrete IL-8 [38–43]. We can speculate that THP-1 monocytes continued to produce IL-8 even if the CPB shear stress stimulus had been removed, that CPB shear stress activated THP-1 stimulated IL-8 release from endothelial cells, or both. Interestingly, IL-8 appears to autocrinally promote its own transcription in monocytes but not in neutrophils [44], suggesting a feed forward loop of self-activation. Future investigation will be needed to determine the source of IL-8 in our system. Apart from the change in IL-8 levels, we observed that the IL-6 level increased significantly at 6 hours of sheared THP-1 cell/endothelial cells coculture. Although studies have linked IL-6 levels to inflammation and postoperative mortality after CPB [45–47], in our model, significant sheared monocyte adhesion was detected at 1 hour into coculture, prior to IL-6 release, and thus suggesting an alternative mechanism. In addition, we demonstrate strong inhibition of CPB-induced THP-1 cell adhesion to endothelial monolayer by the CXCR1/2 inhibitor Reparixin, thus causally implicating the detrimental effect of IL-8 in endothelial function induced by CPB shear stress activated monocytes, which can further lead to vessel leakage and monocytes tissue invasion.

IL-8 is well-known to regulate neutrophil functions by promoting chemiotaxis, causing release of lysosomal enzymes, upregulating of adhesion molecules, increasing intracellular calcium, and priming of the oxidative burst [48–52]. Much less is known on the functions of IL-8 in monocytes. In flow conditions IL-8 stimulates adhesion to E-selectin expressing endothelial cells and promotes monocytes polarization toward an M1 pro-inflammatory phenotype [53–55]. In our system, it appears that IL-8 promoted endothelial specific processes facilitating monocyte adhesion that was strongly inhibited by blocking the IL-8 receptors on endothelial cells. Further studies will be necessary to assess direct effects of IL-8 on CPB shear stress activated monocytes. In addition, endothelial cells have been shown to express the IL-8 receptors CXCR1 and CXCR2 and respond to IL-8 by activating angiogenesis processes, including proliferation, survival, tube morphogenesis and MMP production [56]. The binding of IL-8 on endothelial cells can induce cytoskeletal reorganization through Rho and Rac signaling pathways [57,58], which are correlated to the clustering of E-selectin, intercellular adhesion molecules, and vascular cell adhesion molecules. IL-8 treatment increases the permeability of the endothelium, likely facilitating the subsequent transmigration of monocytes [57,24,59]. Accordingly, our studies also show that IL-8 is a strong inducer of ICAM1 and VCAM1 RNA expression in the microvascular endothelial cells. We also observed IL-8 dependent endothelial cell cytoskeleton rearrangement and VE-cadherin disruption.

Evidence has shown that inflammation leads to organ damage in CPB patients, and that elevated plasma level of IL-8, among other cytokines, positively correlates with acute kidney injury, brain injury in newborns, and intensive care length of stay [23–27]. Experimental evidence also indicates CPB-dependent transmigration and tissue infiltration of leukocytes and neutrophils in particular [60]. In addition, the inflammatory response to CPB can vary by different ages. Pediatric patients are more susceptible to brain injuries than adults after CPB [61]. However, the pathological mechanisms behind organ damage in CPB still remain unresolved. This is the first study to our knowledge to show that CPB shear stress activated monocytes affected endothelial integrity and promoted monocyte adhesion and transmigration in an IL-8-dependent manner, suggesting that IL-8-mediated signaling pathway may play an important role in the onset of CPB-induced inflammatory response and tissue infiltration at the cellular level. These studies further suggest that blocking IL-8 receptors may prove a promising approach to reduce endothelial injury and tissue and organ damage observed in prolong CPB and pediatric patients undergoing CPB.

## Nonstandard Abbreviations and Acronyms

CPB: Cardiopulmonary bypass
IL: Interleukin
TNF-α: Tumor necrosis factor alpha
CXCR: CXC chemokine receptor
G-THP-1: GFP positive THP-1
HNDMVEC: Primary human neonatal dermal microvascular endothelial cell
PVC: Polyvinyl chloride
ICAM1: intercellular adhesion molecule 1
VCAM1: vascular cell adhesion molecule 1
VE-Cadherin: vascular endothelial cadherin

## Acknowledgement

The authors are very thankful to Dr. Nathaniel Peters at UW W. M. Keck Microscopy Center for the necessary training and assistance in acquiring images on the Leica SP8X confocal microscope, which was funded by a S10 OD016240 NIH grant and the UW Student Technology Fee. H. Zhou designed the study, performed all the experiments, analyzed data, and wrote and edited the manuscript. L. N. Tu, N. Vishal helped build the in vitro CPB setup. L. N. Tu, N. Vishal, C. Giachelli, and M. Scatena provided feedback on the manuscript.

